# Abortion and various associated risk factors in dairy cow and sheep in Ili, China

**DOI:** 10.1101/2020.04.20.050872

**Authors:** Xiaoyu Deng, Huan Zhang, Zhiran Shao, Xiaoli Zhao, Qin Yang, Shengnan Song, Zhen Wang, Yong Wang, Yuanzhi Wang, Jinliang Sheng, Chuangfu Chen

**Affiliations:** School of Animal Science and Technology, Shihezi University, 832000-Shihezi City, Xinjiang, China; School of Medicine, Shihezi University, 832000-Shihezi City, Xinjiang, China

## Abstract

We studied livestock abortion and various associated risk factors in the Ili region of northwest China. Livestock abortion prevalence was estimated and correlated with infections (Brucellosis, Salmonellosis, *Mycoplasma* and *Chlamydia* seropositivity) and management (farming type and contact with other herds/flocks) risk factors. The prevalence of cow and sheep abortion induced by *Brucella* was 76.8% (*P*<0.0001) and 84.1% (*P*<0.0001), and *Mycoplasma* caused an estimated 15.5% (*P*=0.025) and 17.6% (*P*<0.001) abortions, respectively. Abortion-related risk factors included mixed farming (cow *P*=0.001, sheep *P*<0.001), contact with other flocks (cow *P*=0.007, sheep *P*=0.003), brucellosis positivity (cow *P*<0.001, sheep *P*<0.001) and *Mycoplasma* positivity (cow *P*=0.031, sheep *P*<0.001). A total of 2996 serum samples (1402 cow, 1594 sheep) were identified by RBPT (Rose Bengal Plate Test), and they showed the seroprevalence of brucellosis in X county was cow 7.1%, sheep 9.1%; in H county was cow 11.7%, sheep 10.7%; and in Q county was cow 4.2%, sheep 9.1%. The seroprevalence of *Mycoplasma* in X county was cow 3.4%, sheep 7.9%; in H county was cow 5.3%, sheep 9.9%; and in Q county was cow 2.1%, sheep 4.3%. A total of 54 samples, including aborted cow (22), sheep (30) fetuses and milk samples (2), were identified as *Brucella melitensis* (*B. melitensis*) positive. A total of 38 *Brucella* were isolated from 16 aborted cow, 20 sheep fetuses and 2 milk samples. All of these isolates were identified, and confirmed, as *B. melitensis*. A phylogenetic tree showed that the *Brucella* isolates closely matched the *B. melitensis* biovar 3 isolated in Inner Mongolia, China, and *B. melitensis* isolated from Norway and India. These results suggest that *B. melitensis* biovar 3 is the main pathogen responsible for cow and sheep abortion and also pose a human health risk. Additionally, livestock reproduction can also be influenced by *Mycoplasma* infection and managerial factors (farming type and contact with other herds/flocks), especially in remote areas.

## Introduction

Ruminants are a major source of meat production in China and are important for food security. Xinjiang Uygur Autonomous Region (XUAR) is located in northwest China and is a major ruminant production province. In 2017, beef production (0.43 million tons) and mutton production (0.58 million tons) in Xinjiang, respectively, accounted for the 6.8% and 12.4% in the total beef and mutton production in China. Ili is located in the western part of XUAR, where the economy is highly dependent on animal production [1, 2]. The combined number of cattle and sheep is approximately 5.76 million in this region. The sheep and cattle are reared under traditional systems, and confined sheep or cattle ranches are the two main feeding systems.

Diseases and poor animal health are major risk factors for animal production in Ili. The viability of sheep and cattle production is largely determined by their reproductive ability, which is influenced by both genetic and environmental factors [3, 4]. Abortion is the most serious threat to livestock, and it is also a public health issue. Abortion is often induced by zoonotic microorganisms [5, 6].

Most ruminants are maintained by poor farmers as way to increase family income. Abortion in sheep and dairy cows has a great impact on the animal production and the health of rural economies [7, 8]. The farming system and communal grazing are often involved in the spread of infectious organisms. There is a need for an improved diagnostics and specific control strategies for maintaining healthy livestock and public health safety [6]. Risk factors responsible for livestock abortion can be classified into infectious and non-infectious [9].

Infectious agents are the main causes of abortion in sheep and cow as compared to non-infectious agents and are generally infectious to humans. The main etiological agents causing sheep and cow abortion are *Brucella, Salmonella, Mycoplasma, Chlamydia abortus* and *Toxoplasma gondii* [9-12].

Ili is an endemic area for brucellosis with high incidences in sheep (4.21%) and cow (6.91%) brucellosis in 2015 (Data from the Center for Animal Disease Control and Prevention of Ili). In this region, most farmers practice mixed farming (both sheep and dairy cows) and use a communal grazing system. Grazing in this environment can expose pregnant animals to pathogens [5, 13].

Recent years, the livestock abortion occurred in an increasing number according to the local Veterinary Department report. However, the causes of these abortions remained unknown. Hence, to protect and sustain the ruminant industry in the Ili region, we need to understand all of the reasons for animal abortion. Thus, the objectives of this study were to: 1) investigate the prevalence of abortion in Ili ruminant flocks and correlated its association with infectious agents (*Brucella, Salmonella, Mycoplasma* and *Chlamydia abortus*) and management (farming type and contact with other flocks) risk factors; 2) isolate and analyze genetic characteristics of the abortion-related pathogens.

## Materials and methods

### Study design

A cross-sectional study was carried out in three counties of the Ili region (X county, H county and Q county) between March and July in both 2017 and 2018. Samples from cows and sheep were collected from smallholder farms. The selection strategies including regions, villages and farms were described in Arif et al [14]. These three counties were selected mostly based on operational convenience, but they also represent a range of agro-ecological zones. Villages within each county were selected randomly by an electronic calculation.

### Herd selection

A total of 325 farms were selected from 25 villages in the three counties, and, given their availability, a maximum of five cows or sheep were randomly sampled in each farm. All of the livestock owners involved were informed about the purpose of this study and provided information about previous vaccinations. The study sampled non-vaccinated animals over two years of age. When there were more than five animals of the required age, five animals were selected randomly from the animals available.

### Sample size

The study population included all the farms in selected villages, but the target population was all of the cows and sheep within the selected villages and all of the villages in selected counties. Several studies reported that *Brucella* is a main pathogen responsible for animal abortion in XUAR [15-17]. Therefore, the sample size was calculated according to the estimated prevalence of brucellosis in these three counties, and the assumed prevalence is listed in Table 1. The minimum samples number of sheep and cows required assumed a closed population, as described previously [18]. The sample size of cows and sheep was estimated to detect a reduction of at least 4% for cows and 6% for sheep brucellosis prevalence with a confidence of 95% and a power of 80% according the following equation [18]:

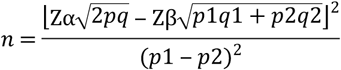

**Table 1.**
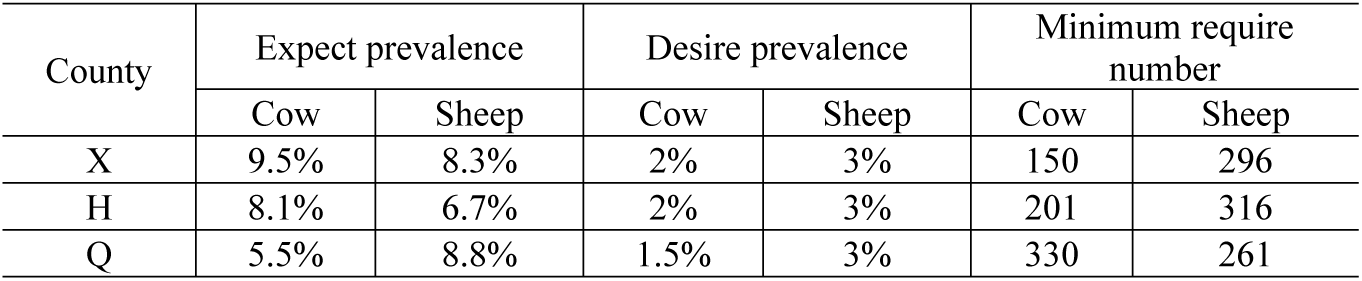
The estimation of minimum livestock number required in this study.

In this equation, *n* is the minimum number of samples required, Z**α** represents the value obtained from standard normal distribution for 95% confidence (1.96), Z**β** represents the value obtained from standard normal distribution for a power of 80% (−0.84), *p*1 represents the estimated prevalence, an expected prevalence for cow and sheep brucellosis in these three counties (listed in Table 1), *p*2 represents the desired brucellosis prevalence for cows and sheep (listed in Table 1), *q*1 is (1-*p*1), *q*2 is (1-*p*2), *p* is (*p*1+*p*2)/2, and *q* is 1-*p*. The minimum number of cows and sheep required in these three counties is shown in Table 1.

### Sample collection

A total of 2996 blood samples (1402 dairy cows and 1594 sheep) were collected from jugular veins using venoject needles (Venoject, China) and stored in 5 mL sterile vacutainer tubes. Additionally, 141 aborted fetuses (66 cow fetuses and 75 sheep fetuses) and 65 milk samples (42 cow milk and 23 ewe milk) were collected. Blood samples were centrifuged at 3000 rpm for 10 min, and the serum was separated into a new sterile tube and stored at −20°C until tested. The milk samples were transported to the laboratory and stored in 4°C. The aborted fetuses were stored at −20°C until processing.

### Laboratory testing

All of the serum samples were screened for antibodies by RBPT, and the positive serum samples were reconfirmed by c-ELISA. Briefly, 30 μL of antigen was mixed with 30 μL of serum on a clean plate. After 3 min, any visible agglutination was considered as positive, and no agglutination was considered as negative. Positive or doubtful samples identified by RBPT were further tested with c-ELISA using the Svanovir *Brucella*-Ab-c-ELISA kits (Svanova Biotech, Uppsala, Sweden) according to the manufacturer’s instructions. The optical density (OD) of each samples were tested twice to obtain the average OD. The cutoff OD of 0.3 was used to identify positive reactions [19]. The sensitivity and specificity of these two methods have been validated as useful tools for brucellosis screening [20, 21]. Additionally, all of the serum samples were screened using the ELISA method to evaluate the changes of *Chlamydia abortus*-specific antibody titer. The mean value of OD was used to identify infected or non-infected livestock [22] and the *Mycoplasma bovis*-specific antibody concentration was determined by *Mycoplasma bovis* MilA IgG ELISA as described previously [23]. The antibodies against *Salmonella spp* were identified using an indirect ELISA kit as described previously [24].

### Risk factors questionnaire

A questionnaire was filled out by participating farm owners. The questionnaire contained information about abortion history in the livestock during previous two years, livestock management risk factors including history of contact with other animals (yes or no) and type of farming; sheep flocks (containing only sheep), cow herds (containing only cows) or mixed groups (containing both sheep and cows).

### PCR examination

Samples of spleen, liver and stomach contents were collected aseptically from aborted fetuses of sheep or cows. The DNA extractions from tissue samples were performed using the TIANamp Genomic DNA Kit (TIANGEN BIOTECH CO., LTD, China) according to the manufacturer’s instructions. Nucleic acid extraction from raw milk was performed as previously described [25]. All of the samples were examined by PCR and the PCR primers used in this study are listed in Table S1 of the Supplementary Material.

### Pathogen isolation

The *Brucella* was isolated from raw milk as previously described [26, 27]. Spleen, liver and stomach contents were crushed and cultured on *Brucella* serum dextrose agar composed of *Brucella* medium base (supplemented with *Brucella* selective antibiotic, OXOID, England) and 5%-10% heat-inactivated horse serum (GIBCO, New Zealand). Plates were incubated with, and without, 5%-10% carbon dioxide at 37°C after inoculation with sample materials. The plates were examined after 3-5 d for bacterial growth. A single clone was chosen for identification. The *Salmonella spp*., *Mycoplasma bovis* and *Chlamydia abortus* were isolated from aborted fetuses or milk samples as described previously [28-30].

### Identification of isolates

The obtained single bacterial clones were identified using PCR targeting the 16S *rRNA* gene [31]. The PCR primers for examination of *Salmonella spp*., *Mycoplasma bovis* and *Chlamydia abortus* are listed in Table S1 of the Supplementary Material. The IS*711* PCR primers were used to identify the species of *Brucella*. PCR products purification and sequencing was conducted as described above. Phylogenetic analysis of isolates was done according to the IS*711* sequence. The sequence distance was determined by the neighbor-joining (NJ) method, and maximum-likelihood algorithms were analyzed using the Molecular Evolutionary Genetics Analysis (MEGA) 7 software [32]. The *Brucella* isolates were characterized by biochemical testing according to the standard strain identification method [33]. The carbon dioxide (CO2) requirement was tested on *Brucella* serum dextrose agar with and without CO2 during the first isolation. Agglutination by A, M and R monospecific antisera was detected by mixing the antisera with the isolate after dilution of the colony. This process was completed at the Center for Disease Prevention and control (CDC) of China in Beijing.

### Statistical analysis

To analyze the risk factors, a preliminary analysis of the data (univariate) was conducted to select the variables with *P* ≤ 0.05 by Chi-square test or Fisher’s exact test. Subsequently, the *P* ≤ 0.05 of variables was analyzed by multivariable logistic regression [34]. The collinearity was verified between each of the independent variables by correlation analysis, and a correlation coefficient >0.9 indicated the variables with strong collinearity. Because of the problem of multicollinearity, one or two variables were excluded from the multiple analysis based on the biological plausibility [35]. Confounding data were evaluated by adding new variables and then monitoring the changes in the model parameters. Large changes (>20%) in the regression coefficients were considered indicative of confounding. The calculations were made using SPSS software 17.0.

## Results

### Distribution of seroprevalence of four abortion-related pathogens in three counties

A total of 2,996 serum samples (1402 dairy cows and 1594 sheep) were collected from X county (cow 352, sheep 353), H county (cow 340, sheep 363) and Q county (cow 710, sheep 878) and then identified by RBPT, c-ELISA and ELISA. The brucellosis positivity for cows and sheep in X county was cow 7.1%, sheep 9.1%; in H county was cow 11.7%, sheep 10.7%; and in Q county was cow 4.2%, sheep 9.1%, which is much higher than the seroprevalence of other pathogens in these three counties (Table 2). However, our results suggest that the *Mycoplasma* infection is an additional threat to livestock reproduction. The *Mycoplasma* positivity for cows and sheep in X county was cow 3.4%, sheep 7.9%; in H county was cow 5.3%, sheep 9.9%; and in Q county was cow 2.1%, sheep 4.3% (Table 2), and its abortion rate for cows and sheep was 15.5% (7/45, *P*=0.025) and 17.6% (18/102, *P*<0.001) (Table 3). The salmonellosis and *Chlamydia abortus* seroprevalence for cows and sheep in these three counties are shown in Table 2.

**Table 2.** Seroprevalence of brucellosis, salmonellosis, *Mycoplasma* and *Chlamydia abortus* in three counties.

| Variables | X County | H County | Q County |
| --- | --- | --- | --- |
| Brucellosis positivity |  |  |  |
| Cow | 25/352 (7.1%) | 40/340 (11.7%) | 30/710 (4.2%) |
| Sheep | 32/353 (9.1%) | 39/363 (10.7%) | 80/878 (9.1%) |
| Salmonellosis positivity |  |  |  |
| Cow | 3/352 (0.85%) | 4/340 (1.2%) | 3/710 (0.42%) |
| Sheep | 6/353 (1.7%) | 6/363 (1.7%) | 4/878 (0.45%) |
| *Mycoplasma* positivity |  |  |  |
| Cow | 12/352 (3.4%) | 18/340 (5.3%) | 15/710 (2.1%) |
| Sheep | 28/353 (7.9%) | 36/363 (9.9%) | 38/878 (4.3%) |
| *Chlamydia abortus*<br>positivity |  |  |  |
| Cow | 1/352 (0.28%) | 1/340 (0.29%) | 3/710 (0.42%) |
| Sheep | 2/353 (0.57%) | 1/363 (0.28%) | 4/878 (0.46%) |

**Table 3.**
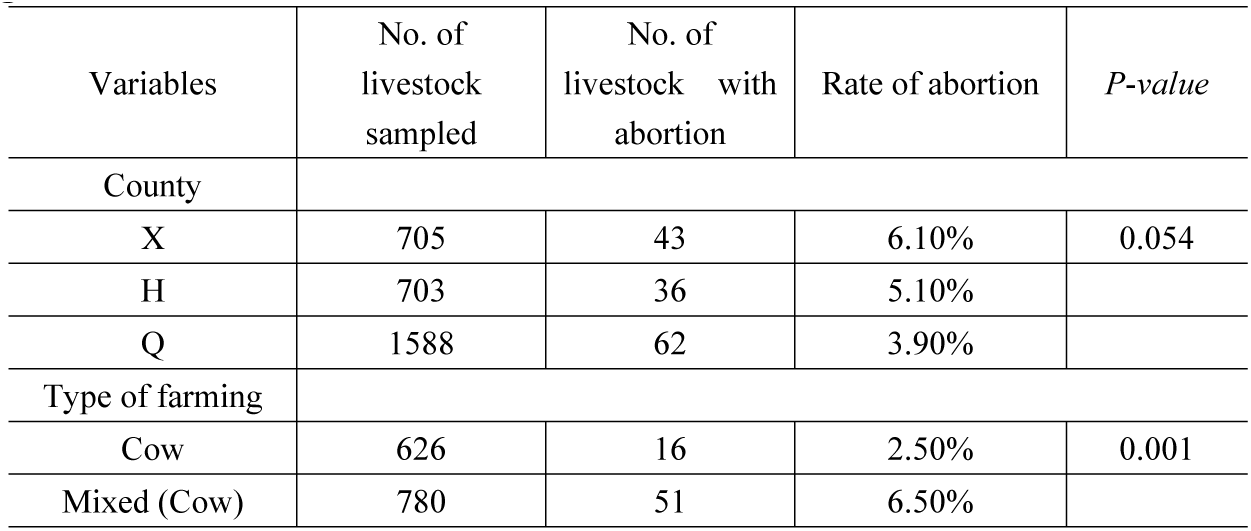
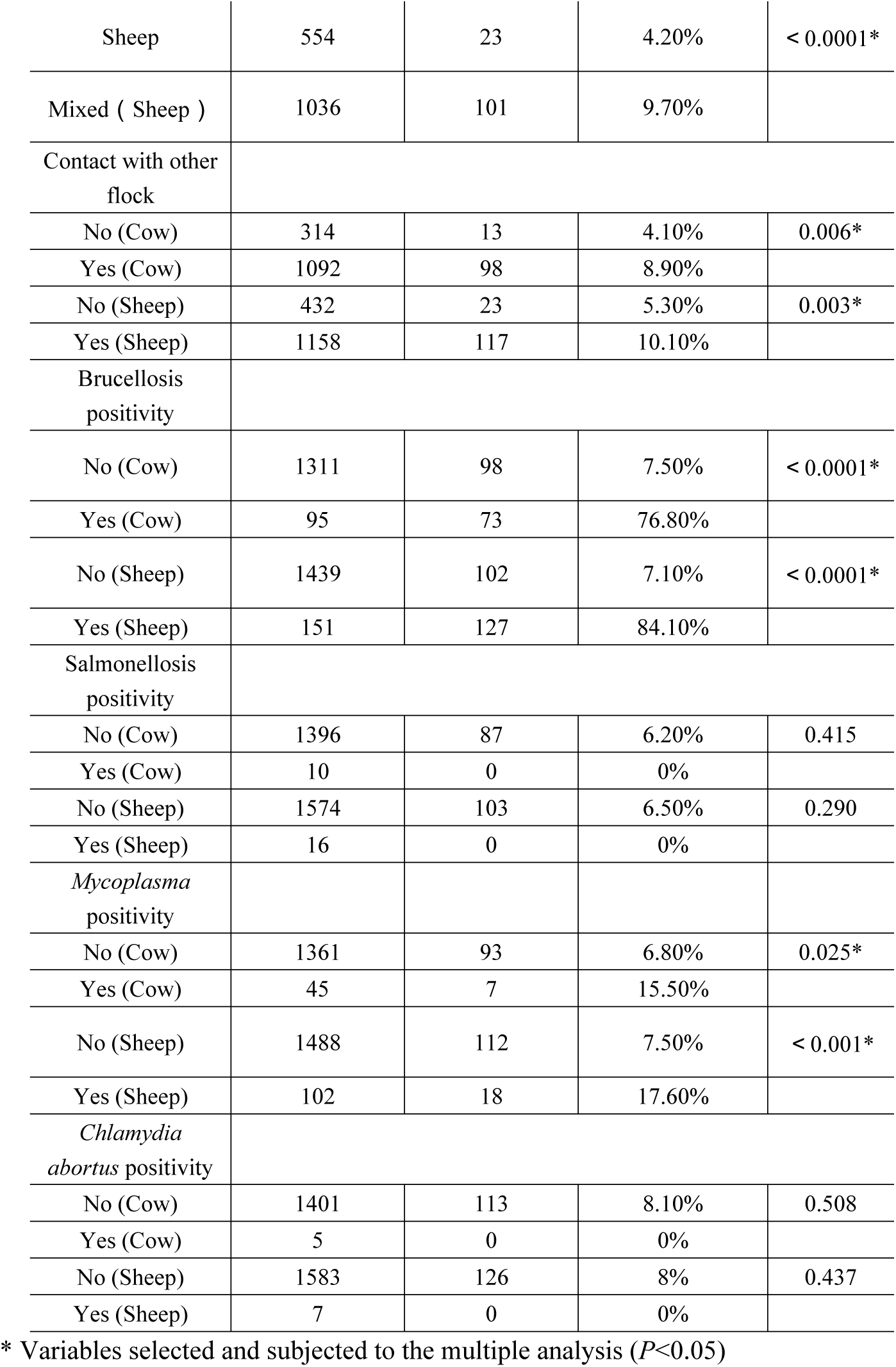
Univariable analysis of abortion-related factors of livestock in the Ili region.

| Variables | No. of livestock sampled | No. of livestock with abortion | Rate of abortion | *P-value* |
| --- | --- | --- | --- | --- |
| County |  |  |  |  |
| X | 705 | 43 | 6.10% | 0.054 |
| H | 703 | 36 | 5.10% |  |
| Q | 1588 | 62 | 3.90% |  |
| Type of farming |  |  |  |  |
| Cow | 626 | 16 | 2.50% | 0.001 |
| Mixed (Cow) | 780 | 51 | 6.50% |  |
| Sheep | 554 | 23 | 4.20% | < 0.0001* |
| Mixed ( Sheep ) | 1036 | 101 | 9.70% |  |
| Contact with other flock |  |  |  |  |
| No (Cow) | 314 | 13 | 4.10% | 0.006* |
| Yes (Cow) | 1092 | 98 | 8.90% |  |
| No (Sheep) | 432 | 23 | 5.30% | 0.003* |
| Yes (Sheep) | 1158 | 117 | 10.10% |  |
| Brucellosis positivity |  |  |  |  |
| No (Cow) | 1311 | 98 | 7.50% | < 0.0001* |
| Yes (Cow) | 95 | 73 | 76.80% |  |
| No (Sheep) | 1439 | 102 | 7.10% | < 0.0001* |
| Yes (Sheep) | 151 | 127 | 84.10% |  |
| Salmonellosis positivity |  |  |  |  |
| No (Cow) | 1396 | 87 | 6.20% | 0.415 |
| Yes (Cow) | 10 | 0 | 0% |  |
| No (Sheep) | 1574 | 103 | 6.50% | 0.290 |
| Yes (Sheep) | 16 | 0 | 0% |  |
| *Mycoplasma* positivity |  |  |  |  |
| No (Cow) | 1361 | 93 | 6.80% | 0.025* |
| Yes (Cow) | 45 | 7 | 15.50% |  |
| No (Sheep) | 1488 | 112 | 7.50% | < 0.001* |
| Yes (Sheep) | 102 | 18 | 17.60% |  |
| *Chlamydia abortus* positivity |  |  |  |  |
| No (Cow) | 1401 | 113 | 8.10% | 0.508 |
| Yes (Cow) | 5 | 0 | 0% |  |
| No (Sheep) | 1583 | 126 | 8% | 0.437 |
| Yes (Sheep) | 7 | 0 | 0% |  |

\* Variables selected and subjected to the multiple analysis ( $P<0.05$ )

### Other livestock management factors involved in abortion

Univariable analysis of abortion-related risk factors (Table 3) found no significant differences among the studied counties (*P*=0.054); the abortion rate in the three regions ranged from 3.9% to 6.1%. However, the management factors were significantly correlated with sheep or cow abortion including the type of farming (cow *P* =0.001, sheep *P*<0.0001) and contact with other herds or flocks (cow *P* =0.006, sheep *P*=0.003). Among the four pathogens, *Brucella* was the main reason for cow or sheep abortion, and the abortion rate of cow or sheep brucellosis was 76.8% and 84.1% (*P*<0.0001) (Table 3). *Mycoplasma* infection also posed a threat to cow and sheep reproduction, and the abortion rates were, respectively 15.5% (*P*=0.025) and 17.6% (*P*<0.001) (Table 3).

### Brucellosis is the main factor responsible for cow and sheep abortion

The abortion-related risk factors analyzed through multivariable logistic regression showed that brucellosis was the biggest risk factor for livestock abortion (Table 4). Our results also showed the brucellosis positivity was significantly associated with cow (*P*<0.0001) and sheep (*P*<0.0001) abortion in the Ili region, and its abortion rates for cow and sheep, respectively, were 76.8% (73/95) and 84.1% (127/151) (Table 3). The Exp (B) values of brucellosis for cow and sheep, respectively, were 41.003 [95% CI 24.396-68.916, *P*<0.001] and 69.362 [95% CI 42.900-112.146, *P*<0.001] times higher than other abortion-related factors including mixed farming, contact with other flocks and *Mycoplasma* infection (Table 4).

**Table 4.** Abortion-related risk factors of livestock in the Ili region.

| Risk factors | Logistic regression coefficient | Standard error | Wald | Exp(B) | 95% CI | P-value |
| --- | --- | --- | --- | --- | --- | --- |
| Mixed farming |  |  |  |  |  |  |
| Cow | 0.978 | 0.292 | 11.230 | 2.658 | 1.501-4.709 | 0.001 |
| Sheep | 0.912 | 0.237 | 14.825 | 2.494 | 1.566-3.971 | <0.001 |
| Contact with other flock |  |  |  |  |  |  |
| Cow | 0.817 | 0.302 | 7.328 | 2.268 | 1.253-4.102 | 0.007 |
| Sheep | 0.691 | 0.235 | 8.650 | 1.999 | 1.260-3.171 | 0.003 |
| Brucellosis positivity |  |  |  |  |  |  |
| Cow | 3.71 | 0.265 | 196.503 | 41.003 | 24.396-68.916 | <0.001 |
| Sheep | 4.237 | 0.245 | 299.073 | 69.362 | 42.900-112.146 | <0.001 |
| *Mycoplasma* positivity |  |  |  |  |  |  |
| Cow | 0.919 | 0.425 | 4.677 | 2.508 | 1.090-5.769 | 0.031 |
| Sheep | 0.969 | 0.278 | 12.151 | 2.633 | 1.528-4.537 | <0.001 |

Exp(B) represent the Odds ratio.

### Molecular detection

In the present study, all of the 75 aborted sheep fetuses, 66 aborted cow fetuses, 42 milk and 23 ewe milk samples were screened by PCR targeting the 16s *rRNA* gene. A total of 54 samples (22 aborted cow fetuses, 30 aborted sheep fetuses, 1 milk and 1 ewe’s milk) were positive and were further identified as *B. melitensis* by targeting the IS*711* gene (data not shown). However, all of these samples were negative for *Salmonella spp*., *Mycoplasma bovis* and *Chlamydia abortus* identified by PCR (data not shown). The nucleotide sequences from this study have been deposited in the GeneBank database (IS*711*: MK913893-MK913898).

### Identification of isolates

A total of 38 (70.37%) *Brucella* isolates were isolated from 54 positive samples, including 20 aborted sheep fetuses, 16 aborted cow fetuses and 1 milk sample and 1 ewe’s milk sample (Table 5). All of the isolates were positive for 16S *rRNA*. The *Brucella* differentiation was performed by PCR utilizing primers specific to the IS*711* gene of *B. melitensis. B. melitensis*-specific DNA fragments with 731 bp were amplified from all isolates, and no DNA was observed in negative control samples. Only part of results is presented in Fig S1 in Supplementary Material. Furthermore. All of the isolates were identified as *B. melitensis* biovar 3 by biochemical testing. The growth of all the 6 isolates on a medium with thionin at 40 µg/mL (1:25000) concentration and basic fuchsin at all concentrations suggested that these isolates were *B. melitensis* biovar 3. Only part of results is presented in Table 6. No *Salmonella spp*., *Mycoplasma bovis* and *Chlamydia abortus* were isolated from aborted fetuses and milk samples.

**Table 5.**
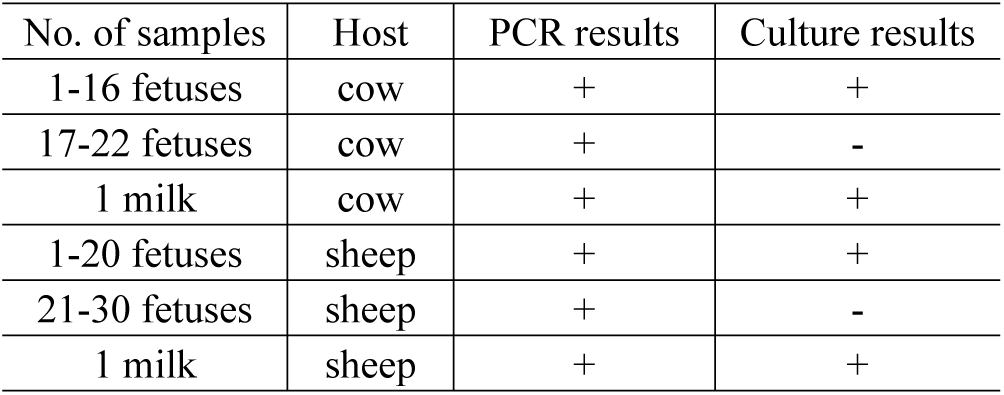
Comparison of PCR and culture results from aborted cow, sheep fetuses and milk samples.

| No. of samples | Host | PCR results | Culture results |
| --- | --- | --- | --- |
| 1-16 fetuses | cow | + | + |
| 17-22 fetuses | cow | + | - |
| 1 milk | cow | + | + |
| 1-20 fetuses | sheep | + | + |
| 21-30 fetuses | sheep | + | - |
| 1 milk | sheep | + | + |

**Table 6.**
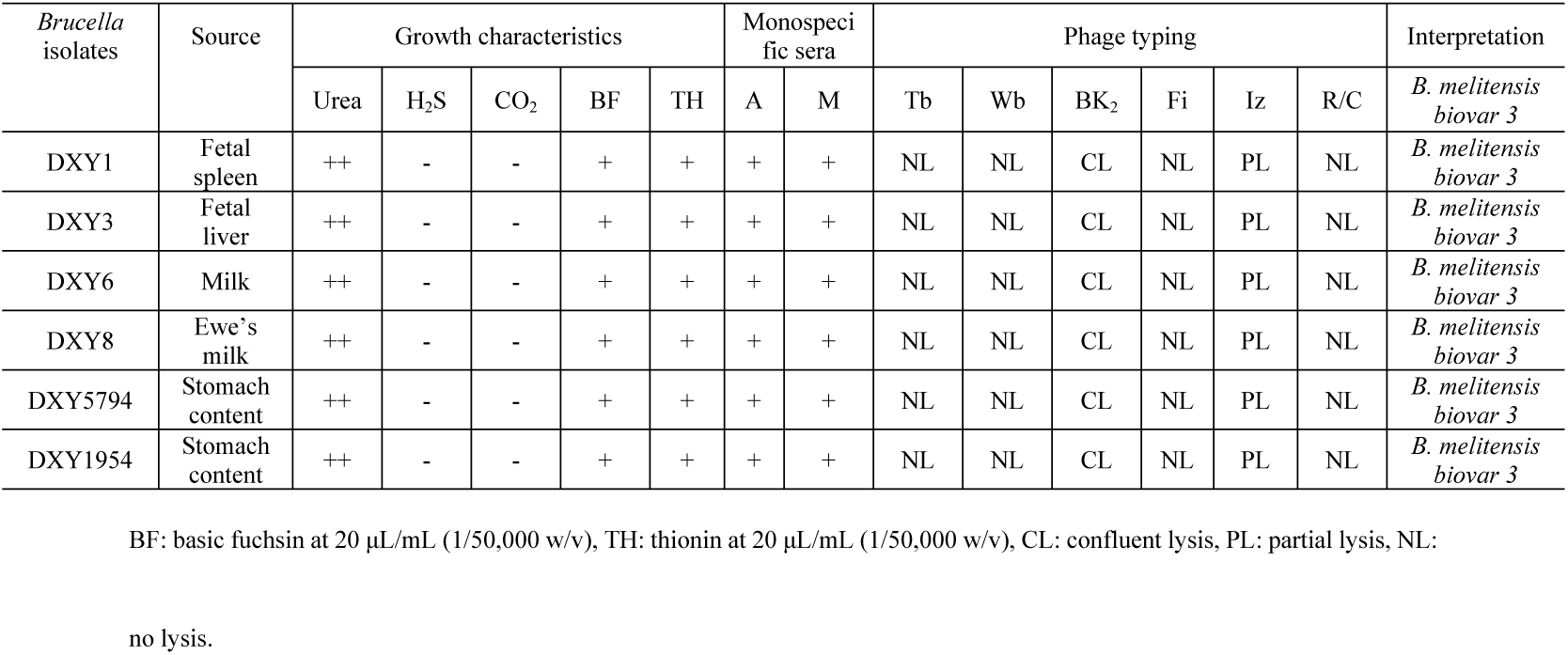
Species and biovar differentiation of the *Brucella* isolates.

| *Brucella*<br>isolates | Source | Growth characteristics |  |  |  |  | Monospecific sera |  | Phage typing |  |  |  |  |  | Interpretation |
| --- | --- | --- | --- | --- | --- | --- | --- | --- | --- | --- | --- | --- | --- | --- | --- |
|  |  | Urea | H<sub>2</sub>S | CO<sub>2</sub> | BF | TH | A | M | Tb | Wb | BK<sub>2</sub> | Fi | Iz | R/C |  |
| DXY1 | Fetal spleen | ++ | - | - | + | + | + | + | NL | NL | CL | NL | PL | NL | *B. melitensis*<br>biovar 3 |
| DXY3 | Fetal liver | ++ | - | - | + | + | + | + | NL | NL | CL | NL | PL | NL | *B. melitensis*<br>biovar 3 |
| DXY6 | Milk | ++ | - | - | + | + | + | + | NL | NL | CL | NL | PL | NL | *B. melitensis*<br>biovar 3 |
| DXY8 | Ewe's milk | ++ | - | - | + | + | + | + | NL | NL | CL | NL | PL | NL | *B. melitensis*<br>biovar 3 |
| DXY5794 | Stomach content | ++ | - | - | + | + | + | + | NL | NL | CL | NL | PL | NL | *B. melitensis*<br>biovar 3 |
| DXY1954 | Stomach content | ++ | - | - | + | + | + | + | NL | NL | CL | NL | PL | NL | *B. melitensis*<br>biovar 3 |

BF: basic fuchsin at 20 µL/mL (1/50,000 w/v), TH: thionin at 20 µL/mL (1/50,000 w/v), CL: confluent lysis, PL: partial lysis, NL:
no lysis.

### Phylogenetic analysis

A phylogenetic tree was constructed based on the 731 bp sequence of the IS*711* repetitive element for all isolates. After sequencing, we found that IS*711* gene sequences from all of these isolates showed 100% similarity (731/731bp). Phylogenetic analysis showed that the *Brucella* isolates closely matched those of *B. melitensis* biovar 3 isolated from cattle in Inner Mongolia, China. Isolates from Norway and India also showed 100% similarity to the isolates of the present study in clade 1 (Fig 1). The isolates of *B. melitensis* from other countries were placed into different clads based on low similarity to the *Brucella* isolates from this study (Fig 1).

**Fig 1.**
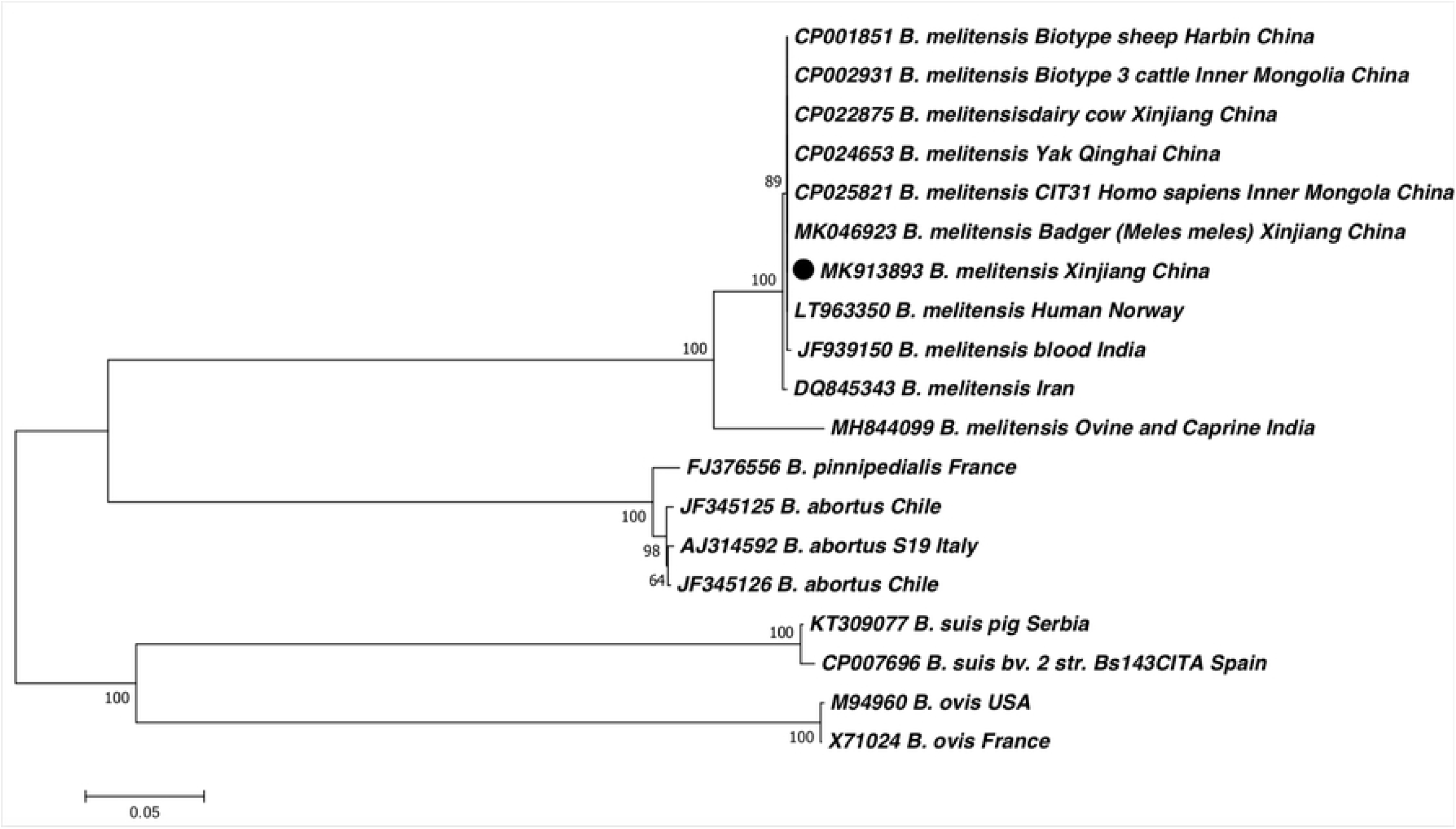
Phylogenetic tree of the IS*711* concatenated sequence of *Brucella melitensis* (•) isolated from aborted cow or sheep fetuses in this study and reference sequences from *Brucella melitensis* retrieved from the GenBank database. The tree was constructed according to the neighbor-joining (NJ; 500 bootstrap replicates) and maximum–likelihood (ML, 1000 bootstrap replicates) analyses using MEGA7. The scale bar represents the inferred substitutions per nucleotide site.

## Discussion

The livestock industry of the XUAR is a major source of its economic growth especially in some remote areas like Ili. However, there are few studies on the prevalence of brucellosis in this region. It has been reported that the brucellosis seroprevalence for cows and sheep in Ili region was cow 1.72%, sheep 1.95% in 2014 [36]. According to the data released from the Center for Animal Disease Control and Prevention of Ili in 2015, the brucellosis seroprevalence for cows and sheep were 6.91% and 4.21%. In the present study, we investigated the seroprevalence of abortion-related pathogens (*Brucella, Salmonella, Mycoplasma* and *Chlamydia abortus*) in three counties (X, H and Q). A total of 2996 cow and sheep serum samples were screened by RBPT, c-ELISA and ELISA. The resulting data showed that the brucellosis was widely prevalent in livestock in all of the studied counties. The seroprevalence for cows and sheep in X county was cow 7.1%, sheep 9.1%; in H county was cow 11.7%, sheep 10.7%; and in Q county was cow 4.2%, sheep 9.1% (Table 2). These data suggest that the disease is distributed within all of the Ili region and can potentially infect all of the susceptible livestock in this region. The results showed that *Mycoplasma* infection was also can influence the livestock reproduction, although its seroprevalence was not as high as brucellosis. The seroprevalence for *Mycoplasma* infection in X county was cow 3.4%, sheep 7.9%; in H county was cow 5.3%, sheep 9.9%; and in Q county was cow 2.1%, sheep 4.3% (Table 2). The abortion rates of *Mycoplasma* positivity for cow and sheep were, respectively, 15.5% (7/45) and 17.6 (18/102) (Table 3). However, Wenhao Ni et al., [37] found that in 2018, the seroprevalence rates of *Mycoplasma* for Hazake sheep and Suffolk sheep were 22.2% and 8.3% in the Ili region. These data are similar to our results except that the higher seroprevalence in Hazake sheep may a breed-related difference.

Many reasons could induce abortion in pregnant animals include infectious factors and non-infectious factors, in which infectious factors include *Brucella, Salmonella spp*., *Mycoplasma bovis, Chlamydia abortus* and *Listeria monocytogenes* [10-12] and non-infectious factors involve heat stress, production stress, seasonal effect, chromosomal and single gene disorders [38-41]. We previously have found that the *B. melitensis* biovar 3 was the main cause of cow and sheep abortion in Nilka county (neighboring X county) in 2016 [42]. However, we could not rule out aborted fetuses caused by non-infectious factors in this study, because part of the fetuses was negative both for those abortion-related pathogens identified by PCR and culture results. In addition to the effects of pathogens on livestock abortion, we found that livestock abortion can also be influenced by livestock management systems including herd and flock size, mixed farming, grazing system and contact with other animals [43, 44]. We used univariable to study management risk factors related to livestock abortion in the Ili region and found statistically significant links with the type of farming (cow *P*=0.001, sheep *P*<0.0001) and contact with other herds/flocks (cow *P*=0.006, sheep *P*=0.003) (Table 3). This may be because these two management factors are easily overlooked by livestock owners.

Bacteria isolation is the gold standard for the diagnosis of brucellosis. We isolated a total of 38 *B. melitensis* biovar 3 isolates from 16 aborted cow fetuses, 20 aborted sheep fetuses and 1 milk and 1 ewe’s milk sample (Table 5). However, there are 16 aborted fetuses that were positive for PCR but negative for culture probably occurred because contamination decreased the rate of *Brucella* isolation. We also identified the *Salmonella spp*., *Mycoplasma* and *Chlamydia abortus* through PCR. But, no aborted fetuses were positive for these pathogens. These results show that *B. melitensis* biovar 3 is the dominant pathogen responsible for sheep and cow abortion.

RBPT and c-ELSA were combined to screen and diagnose brucellosis in China especially in some remote areas. The sensitivity and specificity of these two methods has been described previously [21, 45]. However, these two methods are not good tools for diagnosing brucellosis in the laboratory. We consider that the best way is bacteria isolation and identification. Molecular approaches appeared to be faster and more sensitive than traditional bacteriological tests [46, 47]. The 16S *rRNA* component of the 30S small subunit of prokaryotic ribosomes contains hyper-variable regions that provide species-specific signature sequences useful for bacterial identification. Therefore, the 16S *rRNA* gene can be used as the diagnostic target in the PCR for confirmatory identification of *B. melitensis* [48]. Several studies have demonstrated that the 16S *rRNA* can be used as a rapid tool for *Brucella* identification [48, 49]. In this study, we identified 38 *Brucella* isolates with PCR by targeting the 16S *rRNA* gene in the first round of screening and further identified as *B. melitensis* by the presence of the IS*711* gene. The advantage of this method is that results can be obtained within one d as compared to seven d using traditional microbiological testing.

Brucellosis is principally an animal disease, but >500,000 human cases are reported each year globally [50]. Transmission to humans occurs primarily through contact with infected animals and consumption of contaminated food such as raw milk and its byproducts [51]. This study discovered *B. melitensis* biovar 3 isolates in raw milk and ewe’s milk. This result suggests that *B. melitensis* infection in cows and ewes is a public health issue in China. Infected cows and ewes, as disease reservoirs, can spread contaminated milk to the local human population. We recommend: i) increasing the regular quarantine of brucellosis and timely elimination of infected ewes or cows and their products and ii) implementing a vaccination program for livestock and iii) reducing mixed farming and avoiding contact with other herds/flocks and encouraging livestock owners to learn and adopt new management skills.

## Conclusions

*B. melitensis* biovar 3 was identified as the main pathogen responsible for cow and sheep abortion. *Mycoplasma* infection, mixed farming and contact with other herds and flocks are strongly correlated with livestock abortion. An effective vaccination and control program is advocated for livestock owners in the Ili region to prevent the spread of brucellosis and *Mycoplasma* infection.

## Conflict of interest statement

None of the authors of this paper have a financial or personal relationship with other people or organizations that could inappropriately influence or bias the content of the paper.

## Acknowledgments

We would like to thank LetPub (www.letpub.com) for providing linguistic assistance during the preparation of this manuscript and Shengnan Song for helping construct the phylogenetic tree.

## Supporting information

**Table S1 Primers used in this study**

**Fig S1 PCR product of IS*711* gene amplication**.

